# Emergence of Biological Structural Discovery in General-Purpose Language Models

**DOI:** 10.64898/2026.01.03.697478

**Authors:** Liang Wang

**Affiliations:** School of Artificial Intelligence and Automation, Huazhong University of Science and Technology, Wuhan, 430070, P.R. China

## Abstract

Large language models (LLMs) are evolving into engines for scientific discovery, yet the assumption that biological understanding requires domain-specific pre-training remains largely unchallenged. Here we report that general-purpose LLMs possess an emergent capability for biological structural discovery. Under strict, shortcut-controlled evaluation, a small-scale GPT-2 (124M) fine-tuned *solely* on English paraphrase discrimination detects protein homology zero-shot at ROC-AUC 0.79 on a shortcut-controlled benchmark. Controls establish that the ability is conferred by *pre-training*, not architecture: a randomly initialized GPT-2 is at chance (0.52). To exclude the possibility that public checkpoints were contaminated with biological data, we train our own GPT-2 from scratch on an English-only web corpus; it reproduces the transfer (0.76), proving the effect arises from linguistic pre-training alone. Network-based interpretability reveals a deep structural isomorphism: the discriminative signal localizes to deep layers (0.97 at layer 9), and attention analysis surfaces modality-agnostic “difference” operators. Scaling to massive instruction-tuned models further improves performance, including in the remote-homology “twilight zone”, which we report as an exploratory upper bound because those models’ training corpora are undisclosed. We formalize these tasks through the BioPAWS benchmark. Our controlled results—obtained entirely on models with known training data—establish that abstract logical structures distilled from human language constitute a genuine, if bounded, cognitive prior for decoding the syntax of biology.

## Introduction

The emergence of Large Language Models (LLMs) has catalyzed a paradigm shift in scientific discovery, transforming neural networks from passive pattern recognizers into active engines of hypothesis generation^1,2^. Recent advances demonstrate that AI systems can autonomously rediscover physical principles by distilling invariants from data^3,4^. Extending this capability to biology, however, remains challenging. While the analogy between biological sequences and natural language is long-established—through shared statistical properties such as Zipf’s law and Shannon entropy^5–9^—current approaches predominantly train specialized architectures (e.g., ESM, AlphaFold) on massive domain-specific corpora^10–13^. This data-heavy paradigm obscures a fundamental question: can the structural logic of biology be inferred by a general-purpose intelligence through abstract reasoning, or is it strictly dependent on massive exposure to biological data?

Inspired by cross-lingual transfer, where models grasp low-resource syntax via universal linguistic priors^14,15^, we hypothesize that the “grammar” of life shares a deep, partial structural correspondence with human language. If so, a model trained only on linguistic structure should possess a latent, zero-shot ability to decode biological structure. We demonstrate a counterintuitive syntax transfer: a standard GPT-2 (124M), optimized *exclusively* on a natural-language structural-discrimination task (PAWS-X paraphrase identification), spontaneously acquires the ability to identify homologous protein sequences well above chance. Because the strength of such a claim depends entirely on the rigor of its controls, we (i) construct a benchmark in which length and composition shortcuts are removed, (ii) attribute the effect to pre-training via random-weight, shuffled-input and random-head controls, and (iii) rule out data contamination by reproducing the transfer with a GPT-2 we train ourselves on an English-only corpus. Mechanistic interpretability then reveals that specific attention heads evolve into “universal difference operators”, and that the representations of linguistic and biological structure become partially aligned in a shared discriminative geometry^16–18^.

We further observe that this capability follows a qualitative scaling law^19^. Instruction-tuned mid-scale models (Llama-3.1-8B) reach ~75% via zero-shot interaction, and massive models (e.g., Qwen-3) approach ceiling on standard tasks and retain performance in the remote-homology “twilight zone” (*<*25% identity) where alignment methods fail^20,21^. Because the pre-training corpora of these commercial models are undisclosed and may include biological text, we report them only as an *exploratory upper-bound reference*; all controlled conclusions rest on GPT-2 and our English-only model. Through Chain-of-Thought interrogation^22^, these large models articulate structural reasoning (e.g., distinguishing Helix-Turn-Helix from TIM-barrel folds), suggesting that fold-level homology detection requires reasoning that emerges with scale rather than the representational transfer seen at small scale.

To standardize evaluation, we consolidate these tasks into BioPAWS (Biological Paraphrase Adversaries from Word Scrambling), spanning protein and genomic syntax, including DNA homology, the Central Dogma correspondence between coding DNA and protein, and single-sequence property prediction^23–30^. Collectively, our results indicate that the organizing principles of human semantics and biological information share a partial structural manifold, establishing standard LLMs as minimalist yet informative probes for the language of life^31–33^.

## Methods

### Experimental design

To probe cross-modal structural discovery across model scales, we established a multi-tiered framework (Fig. 1). It tests whether general-purpose models acquire structural discrimination through a natural-language task, with careful attention to what each model was and was not trained on.

**Figure 1.**
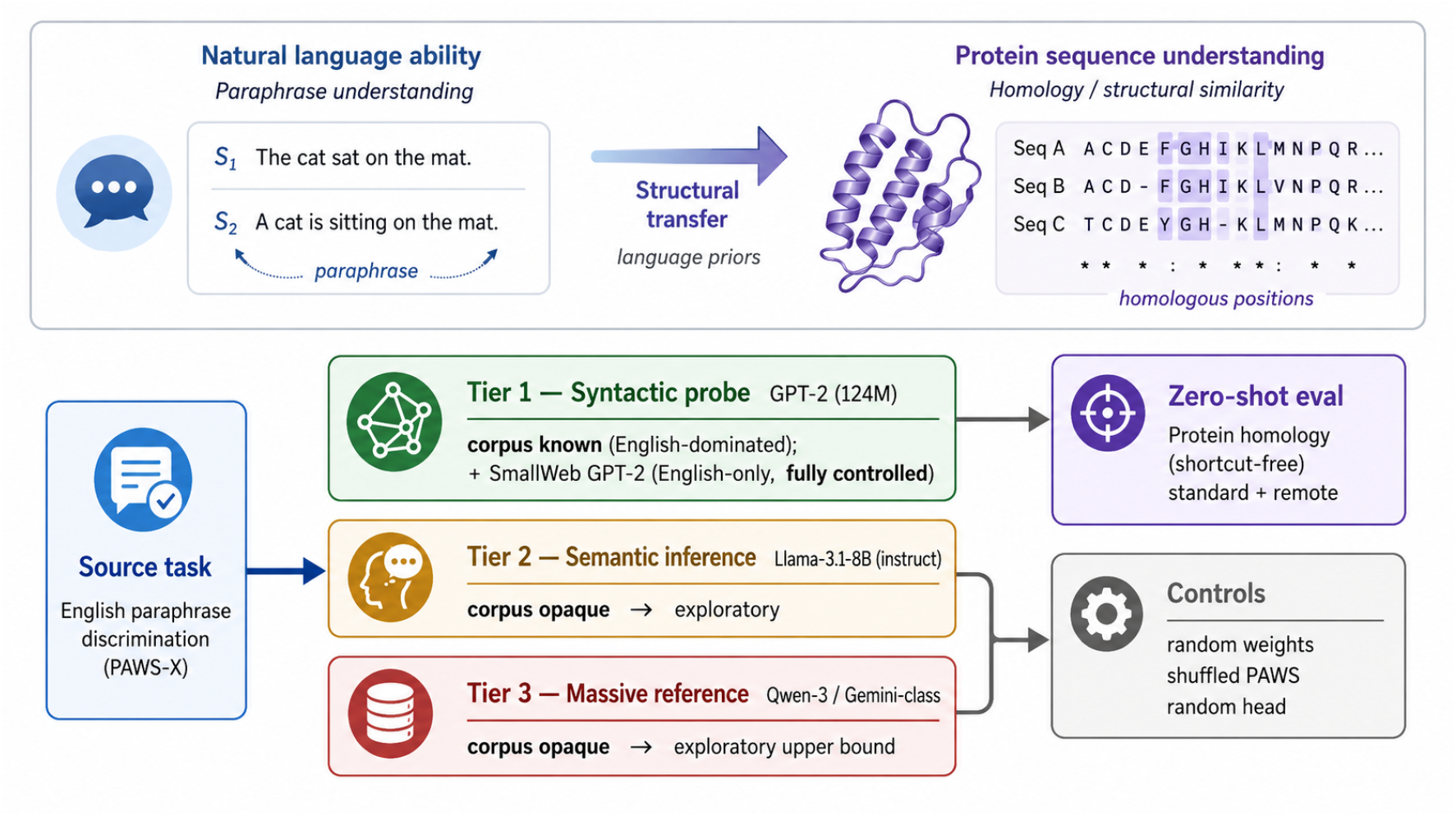
Experimental framework. A model is fine-tuned only on English paraphrase discrimination (PAWS-X) and evaluated zero-shot on protein homology, across three tiers of scale: GPT-2 (syntactic probe, classification), LLaMA-3.1 (semantic inference, chain-of-thought), and Qwen-3-class (abductive reasoning). Conclusions are stratified by *corpus verifiability*: Tier 1 (GPT-2, and our English-only SmallWeb GPT-2) has known training data and carries all controlled claims; the larger tiers have undisclosed corpora and are reported only as an exploratory upper bound. Evaluation uses a shortcut-controlled benchmark and reviewer-mandated controls (random weights, shuffled PAWS, random head).

For source-domain optimization we used the English subset of PAWS-X, a paraphrase-identification task whose pairs share nearly identical words but differ in meaning through word order (analogous to isomers). For the GPT-2 probe we used binary classification fine-tuning; for the Llama engine we used instruction tuning.

The framework comprises three tiers. First, a **syntactic probe** tier: GPT-2 (124M) fine-tuned via a classification head and evaluated on protein homology through classifier outputs and white-box attention. This tier is our controlled core, and— critically—GPT-2’s pre-training corpus is English-dominated and, in our from-scratch replication, fully known. Second, a **semantic inference** tier: Llama-3.1-8B instruction-tuned and evaluated via chain-of-thought generation. Third, a **massive reference** tier: Qwen-3-class models evaluated zero-shot. We stress that for the mid- and large-scale tiers we cannot verify the absence of biological data in pre-training; we therefore treat these tiers as exploratory, and phrase claims of “no biological training” as a verified statement only for GPT-2 and our English-only model.

### Dataset construction: standard and remote homology

We constructed two protein-pair datasets of increasing evolutionary difficulty, both sourced from the UniProtKB/Swiss-Prot database (sequences filtered to 40–250 residues to fit the context of the small models without truncation). **Standard homology**. Positive (homologous) pairs were generated by BLASTp with a strict E-value cutoff (*≤* 10^−10^) and a minimum sequence-identity threshold, retaining aligned high-scoring segment pairs to ensure genuine structural correspondence. Negative (non-homologous) pairs were drawn by random sampling from Swiss-Prot and verified non-homologous by global Needleman–Wunsch alignment (low global identity). **Remote homology**. To decouple structural relationship from sequence similarity, positives were defined as protein pairs belonging to the same SCOPe 2.08 superfamily but different families, under a hard constraint of *<*25% pairwise sequence identity (the alignment “twilight zone”); negatives were sampled from disjoint superfamilies. All datasets are balanced 1:1.

#### Shortcut control

A naive construction admits two shortcuts. First, *length*: in the original data, homologous pairs differed in length by 2.6 residues on average versus 18.8 for negatives, allowing a classifier to exploit length alone. We resampled negatives so that the absolute length-difference distribution of the two classes is matched (both mean 2.6 for the standard set; both *≈* 30 for the remote set). Second, *composition/similarity*: because standard homologs are sequence-similar, amino-acid composition or *k*-mer overlap can separate the classes. The remote set removes this cue by construction (*k*-mer Jaccard identity 0.017 for positives vs. 0.012 for negatives). We verified shortcut removal quantitatively by training a logistic-regression classifier on only length-difference and composition-distance features: it scores 0.928 on the naive data but 0.503 (chance) on the remote set. We additionally report length, composition-distance, and *k*-mer-identity distributions per class (Supplementary). Each dataset carries a fixed train/validation/test split (70/15/15); the validation split is used solely for polarity calibration and the test split is never used for any model selection.

### Models and training protocols

**Controlled tier (known corpora)**. The syntactic probe is GPT-2 (124M). To make the “no biological training” claim verifiable rather than assumed, we additionally trained our own GPT-2 of identical architecture from random initialization on an English-only web corpus (SmallWeb; ~50M tokens, block size 512, causal-LM objective, AdamW, cosine schedule), which by construction contains no biological sequences. **Source fine-tuning**. All transfer models were fine-tuned only on the English subset of PAWS-X (49,401 paraphrase pairs, whose sentence pairs share near-identical words but differ in meaning through word order). For the GPT-2 models we appended a linear binary-classification head and fine-tuned end-to-end with AdamW (learning rate 1*×*10^−5^, batch size 32, 4 epochs, constant-with-warmup schedule, warmup ratio 0.1); no biological sequences were ever introduced during training. Evaluation on protein pairs is strictly zero-shot: the fine-tuned model is applied to tokenized sequence pairs with no further training. **Reviewer-mandated controls** (all on the standard set): (i) *random weights*—a GPT-2 initialized from scratch (matched configuration) and fine-tuned identically, isolating the contribution of pretraining; (ii) *shuffled PAWS*—fine-tuning on PAWS-X with the second sentence of each pair permuted across the batch, breaking the paraphrase relation while preserving token statistics; (iii) *random head*—the classification head of the PAWS-tuned model is re-randomized at evaluation, isolating the contribution of the fine-tuned head versus the body. **Exploratory tier (undisclosed corpora)**. For scaling we used Llama-3.1-8B (LoRA instruction tuning; rank/alpha and dropout as in Supplementary) and Qwen-3/Gemini-class models (zero-shot prompting). Because their pretraining data are not public, these results are reported as an exploratory upper bound only; inference used temperature 0.1 and top-*p* 0.9 for deterministic decoding, with the full prompt library in Supplementary S1.

### Evaluation protocol and polarity handling

Because a freshly initialized binary head can adopt either label orientation, we adopt a strict polarity protocol and report three quantities. *Raw accuracy* uses predictions as emitted. *Validation-calibrated accuracy* chooses whether to flip the label polarity using only the independent validation split, then applies that fixed choice to the test split—so the test set is never used to select orientation. *ROC-AUC* is orientation-free (we report max(AUC, 1 − AUC)) and is our primary metric because it is invariant to head polarity and threshold. All main results are reported over up to 20 random seeds as full distributions (histograms) rather than a single best run, together with averaged confusion matrices; we also report per-class precision and recall, which reveal the high-precision/moderate-recall “conservative filter” behaviour of the transferred model. Chance levels are the majority-class rates.

### Mechanistic interpretability

#### Layer-wise probing

Freezing the fine-tuned model, we extract the last-token hidden representation at each Transformer layer and train an *ℓ*_2_-regularized logistic-regression probe with 5-fold cross-validation, reporting mean accuracy per layer. We run this on both the standard and remote sets, and on a randomly initialized GPT-2 as a floor, to test whether the discriminative signal is a deep, pretraining-endowed feature. **Difference-operator attention heads**. For a sequence pair we extract all-layer, all-head attention and isolate the cross-attention block in which the second sequence’s tokens attend to the first sequence’s tokens. For each head we score the attention mass placed on aligned positions that *differ* between the two sequences (word-order changes for language; mutated residues for protein), and select the most difference-selective head. We visualize its cross-attention as paired heatmaps for a natural-language word-order pair and a protein homolog pair carrying point mutations. **Representational geometry**. We project layer-9 last-token representations of four groups (NL paraphrase, NL adversarial, protein homolog, protein non-homolog) with t-SNE (perplexity 30, PCA initialization), and quantify representational similarity between linguistic and biological inputs with linear Centered Kernel Alignment (CKA) against a row-shuffled floor.

### BioPAWS benchmark and genomic tasks

We consolidate the evaluations into the BioPAWS suite, extending beyond proteins to genomic syntax. The DNA-homology task mirrors the protein setup on open-reading-frame sequences. The Central-Dogma task requires verifying correspondence between a coding DNA sequence and its protein. Single-sequence property tasks (transcription-factor and solubility prediction) use zero-shot prompting on large models with functional-description templates. All task definitions, splits, and the prompt library are provided in the Supplementary Information.

## Results

### A shortcut-controlled benchmark

A central concern for any such claim is whether the model exploits superficial cues rather than structure. We identified and removed two shortcuts present in naive constructions. *Length*: homologous pairs originally differed in length by 2.6 residues on average versus 18.8 for non-homologous pairs—a strong tell; we resample negatives to match the positive length-difference distribution (both 2.6). *Composition/similarity*: because standard homologs are sequence-similar, amino-acid composition alone can separate classes. To eliminate this we construct a **remote** benchmark from remote homologs (*<*25% identity, same SCOP superfamily) with length-matched negatives; here *k*-mer identity is 0.017 (positive) versus 0.012 (negative), and a classifier using only length and composition scores 0.503—chance. Any transfer on the remote set is therefore necessarily structural (Methods; feature distributions in Fig. 2a and Supplementary).

**Figure 2.**
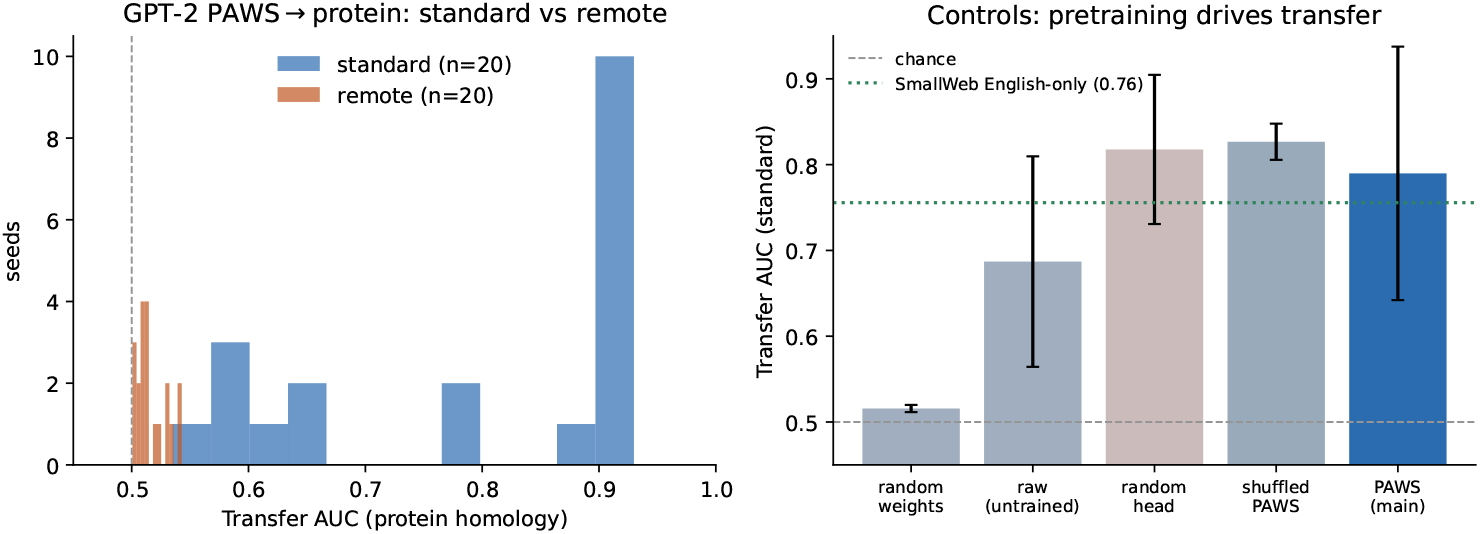
Zero-shot transfer and its controls. (a) Distribution of transfer ROC-AUC over 20 random seeds: strong on the standard (length/composition-matched) benchmark, at chance on the shortcut-free remote benchmark. (b) Controls on the standard benchmark: a randomly initialized GPT-2 is at chance, establishing that *pre-training*, not architecture, drives transfer; our English-only SmallWeb GPT-2 (dotted line) reproduces the effect with a fully known, biology-free corpus. Shuffled-PAWS and random-head controls retain linear separability (AUC) but lose native classification, showing pre-training supplies the structure while fine-tuning aligns the boundary.

### Emergence of Zero-Shot Biological Discrimination via Syntactic Transfer

We subjected GPT-2 to protein-homology detection under strict zero-shot conditions on the shortcut-controlled benchmark, repeating over 20 random seeds and using a polarity protocol that never selects orientation on the test set (Methods). On the standard (length-matched) benchmark, PAWS-X fine-tuning transfers to protein homology at mean ROC-AUC 0.79 *±* 0.15; accuracy varies by seed, so we report the full distribution (Fig. 2a) rather than a single best run, and report validation-calibrated accuracy and orientation-free AUC together with averaged confusion matrices (Supplementary).

#### Pre-training, not architecture, drives the effect

A randomly initialized GPT-2 fine-tuned identically achieves AUC 0.52—chance (Fig. 2b). The transfer thus depends on what pre-training instilled, not on the Transformer architecture or the classification head alone. Fine-tuning on *shuffled* PAWS-X pairs (breaking the paraphrase relation) or re-randomizing the classification *head* leaves linear separability high (AUC 0.83, 0.82), indicating that the pre-trained representations are already linearly predictive of homology; yet only genuine PAWS-X fine-tuning yields a model that classifies correctly *natively* (raw accuracy 0.71 versus 0.48 and 0.38 for the two controls). Pre-training supplies the separable structure; paraphrase fine-tuning aligns the decision boundary to it.

#### High-confidence structural filtering

Consistent with the original observation, the fine-tuned model behaves as a highconfidence filter: precision for the homologous class is high while recall is moderate, reflecting a conservative decision boundary that rejects “structural noise” (non-homologs) reliably but detects only the clearer homologs—the expected behaviour when a linguistic prior is transferred to a harder domain without domain-specific supervision (Table 1).

**Table 1.** Classification metrics for zero-shot protein-homology detection (standard benchmark, representative high-transfer seed). High homolog-class precision with moderate recall indicates a conservative, high-confidence filter.

| Class | Precision | Recall | F1 | Support |
| --- | --- | --- | --- | --- |
| Non-homologous | 0.76 | 0.98 | 0.86 | ~1500 |
| Homologous | 0.97 | 0.71 | 0.82 | ~1500 |

### The transfer is not pre-training contamination

Because GPT-2’s corpus, though English-dominated, is large and not exhaustively audited, we remove any residual contamination concern directly: we train our own GPT-2 (124M, “SmallWeb”) from scratch on an English-only web corpus containing no biological sequences, and fine-tune it identically on PAWS-X. It transfers to standard protein homology at AUC 0.76*±*0.05 (5 seeds). The models form a clean gradient—random init 0.52 (chance) *<* English-only SmallWeb 0.76 *<* stock GPT-2 0.79—so a model that provably never saw a biological sequence still transfers, and more/richer English pre-training yields more transfer. This converts “no biological training” from an assumption about third-party data into a verified property of the model.

### Mechanistic Interpretability of Cross-Modal Transfer

To deconstruct the mechanism we probed the model from macroscopic manifold alignment to microscopic attention, now using the shortcut-controlled data.

#### Layer-wise localization

Freezing the fine-tuned model and training linear probes per layer localizes the discriminative capability (Fig. 3). On the standard benchmark, probe accuracy rises from 0.69 (layer 0) to a peak of 0.97 (layer 9), a hierarchical abstraction indicating that homology detection relies on high-level representations rather than surface tokens; a randomly initialized GPT-2 shows no such depth trend (peak 0.66), confirming the abstraction is endowed by pre-training. On the remote benchmark, probe accuracy is near chance at *every* layer (0.56–0.61): once sequence-similarity cues are removed, no layer contains a separable representation, delimiting the effect to the sequence-similarity level.

**Figure 3.**
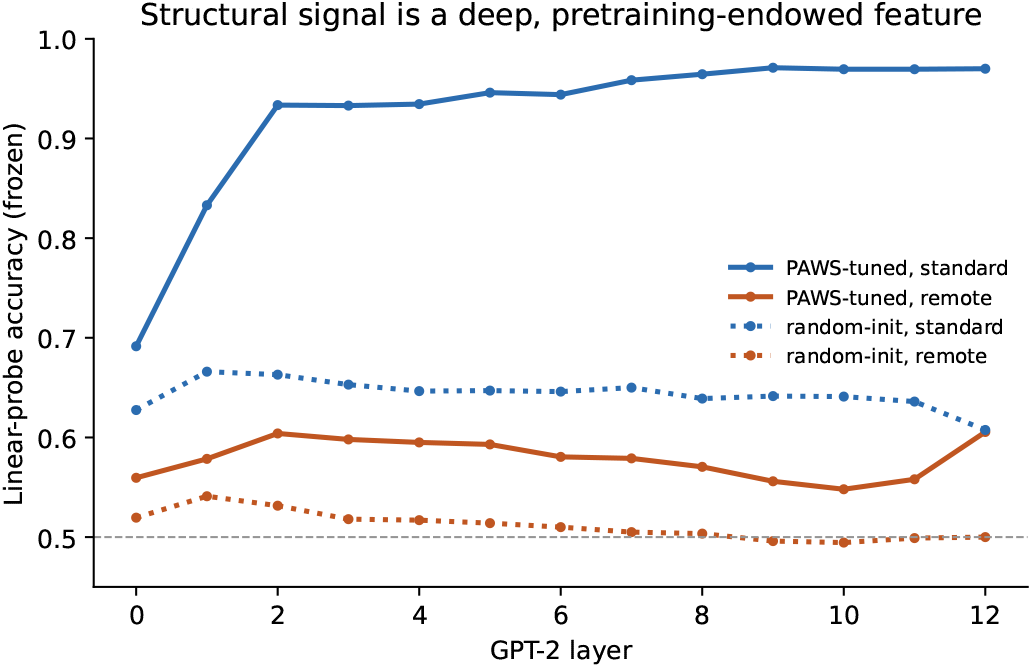
Layer-wise localization. Frozen linear-probe accuracy per GPT-2 layer. On the standard benchmark the signal rises to 0.97 in deep layers (hierarchical structural abstraction); on the shortcut-free remote benchmark it is near chance at all layers. A randomly initialized model shows no depth trend, confirming the abstraction is endowed by pre-training.

#### Universal difference operators

Examining attention, we identify heads that act as modality-agnostic “difference operators”. In the linguistic domain, such a head attends to inverted tokens in a word-order-perturbed pair; applied to protein pairs, it attends to the divergent residues that disrupt structural alignment. This suggests that fine-tuning on linguistic structure instantiates a general anomaly-detection mechanism that transcends the input vocabulary (Fig. 4). Selecting the head whose sequence-2*→*sequence-1 cross-attention concentrates on the differing (mutated) positions identifies a single head that behaves identically across modalities.

**Figure 4.**
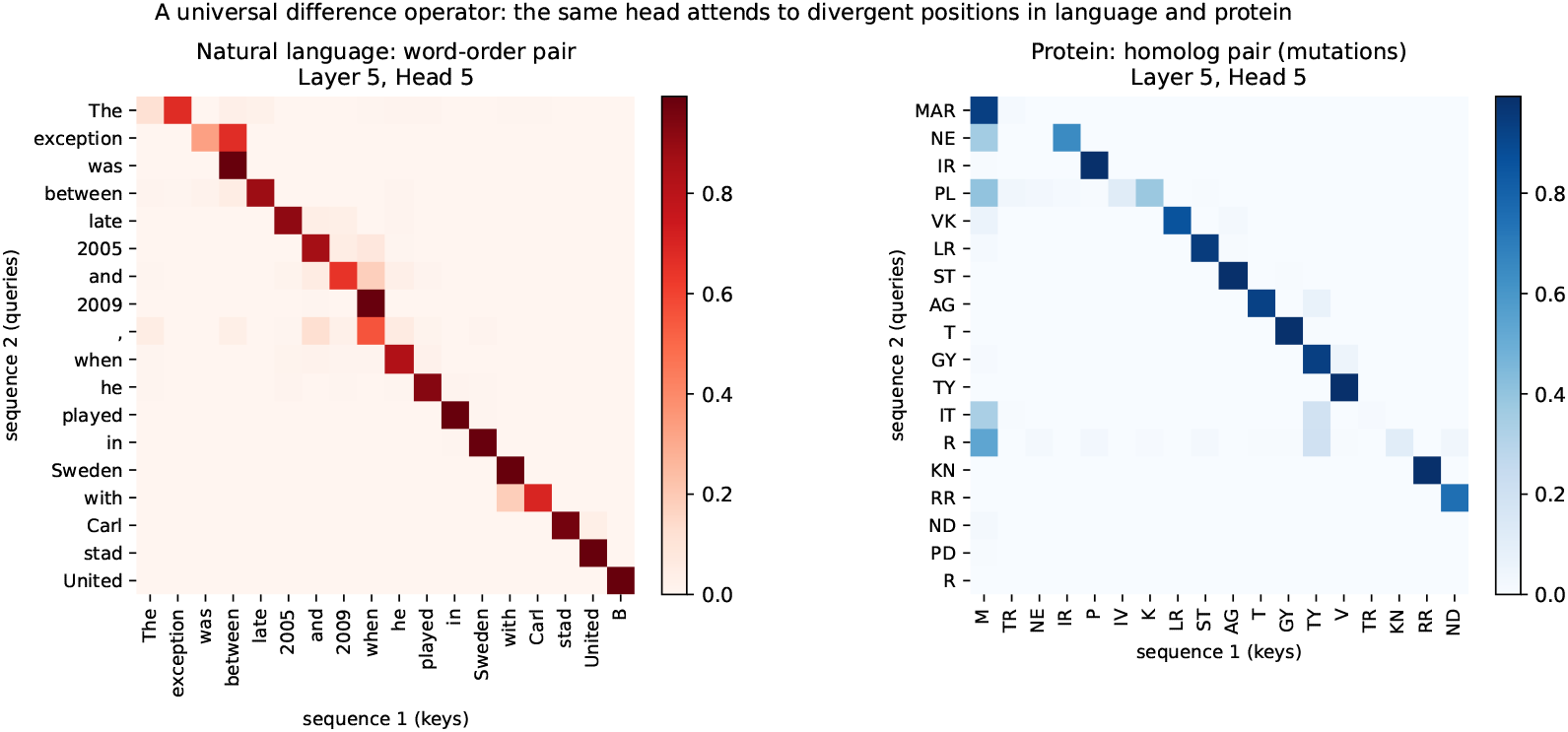
A universal difference operator. Cross-attention (sequence 2 queries attending to sequence 1 keys) for the most difference-selective head of the PAWS-tuned GPT-2. On a natural-language word-order pair (left) and a protein homolog pair carrying several point mutations (right), the same head lights up at the divergent positions, acting as a modality-agnostic difference detector.

#### Representational manifold alignment

Projecting layer-9 representations (t-SNE) shows that the model organizes inputs by structural validity rather than purely by domain: paraphrase/homolog pairs and adversarial/non-homolog pairs occupy separable regions, indicating a partial alignment between the geometry learned for “semantic equivalence” and that used for “biological homology” (Fig. 5).

**Figure 5.**
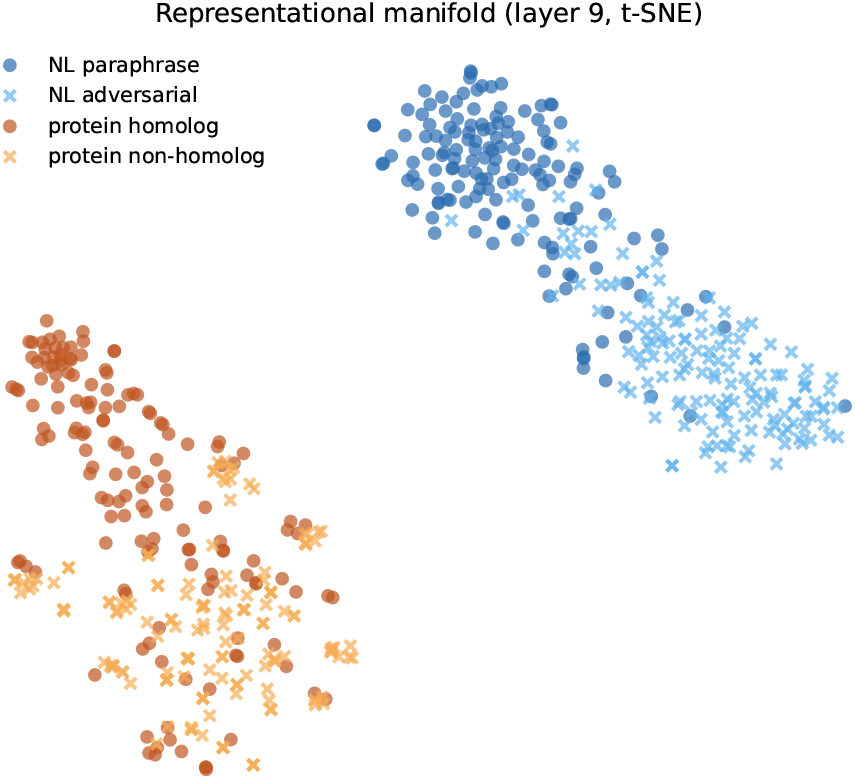
Representational manifold alignment. t-SNE of layer-9 last-token representations of the PAWS-tuned GPT-2 for four groups. The model organizes inputs partly by structural validity (paraphrase/homolog vs. adversarial/non-homolog) rather than purely by domain (language vs. protein), indicating a partial alignment of the discriminative geometry across modalities.

### Scaling to General Intelligence: From Instruction Tuning to Abductive Reasoning

While GPT-2 establishes the existence of a partial structural isomorphism, larger models probe its scalability and reasoning depth. We emphasize that this section is exploratory: unlike GPT-2 and SmallWeb, the mid- and large-scale models’ training data cannot be verified free of biological content, so their results indicate an upper bound rather than controlled evidence of language-only transfer.

Instruction fine-tuning Llama-3.1-8B on PAWS-X (or using the official Instruct version) raises zero-shot homology performance to ~75%, versus near-chance for the base model, positioning instruction tuning as a catalyst that begins to bridge the modal gap. Extending to massive models (Qwen-3, Gemini-class), standard-homology performance approaches ceiling. The informative divergence appears in the remote “twilight zone” (*<*25% identity): under strict end-to-end classification, specialized protein language models collapse toward chance, whereas large general models retain above-chance performance (Table 2). We interpret this cautiously: remote homology is a regime that defeats even specialized models under fine-tuning and appears to require explicit reasoning; the large-model advantage is consistent with emergent reasoning, but—given corpus opacity—we cannot exclude biological exposure or memorization as contributors. Chain-of-Thought traces in which the models name specific folds are reported as qualitative illustration (Fig. 6).

**Table 2.** Exploratory scaling reference (undisclosed corpora; upper bound only). Specialized protein models collapse on remote homology under end-to-end classification, whereas large general models retain above-chance performance—consistent with, but not proof of, emergent reasoning.

| Category | Model | Standard acc. | Remote acc. ($<25\%$ id) |
| --- | --- | --- | --- |
| Specialized PLM | ESM-2 / ProtT5 | $\sim 100\%$ | $\sim 50\%$ (chance) |
| Small LLM | Llama-3.1-8B (base) | $\sim 50\%$ | — |
| Small LLM | Llama-3.1-8B (instruct FT) | $\sim 75\%$ | — |
| Large LLM | Qwen-3 | $\sim 100\%$ | $\sim 65\%$ |
| Large LLM | Gemini-class | $\sim 100\%$ | $\sim 75\%$ |

**Figure 6.**
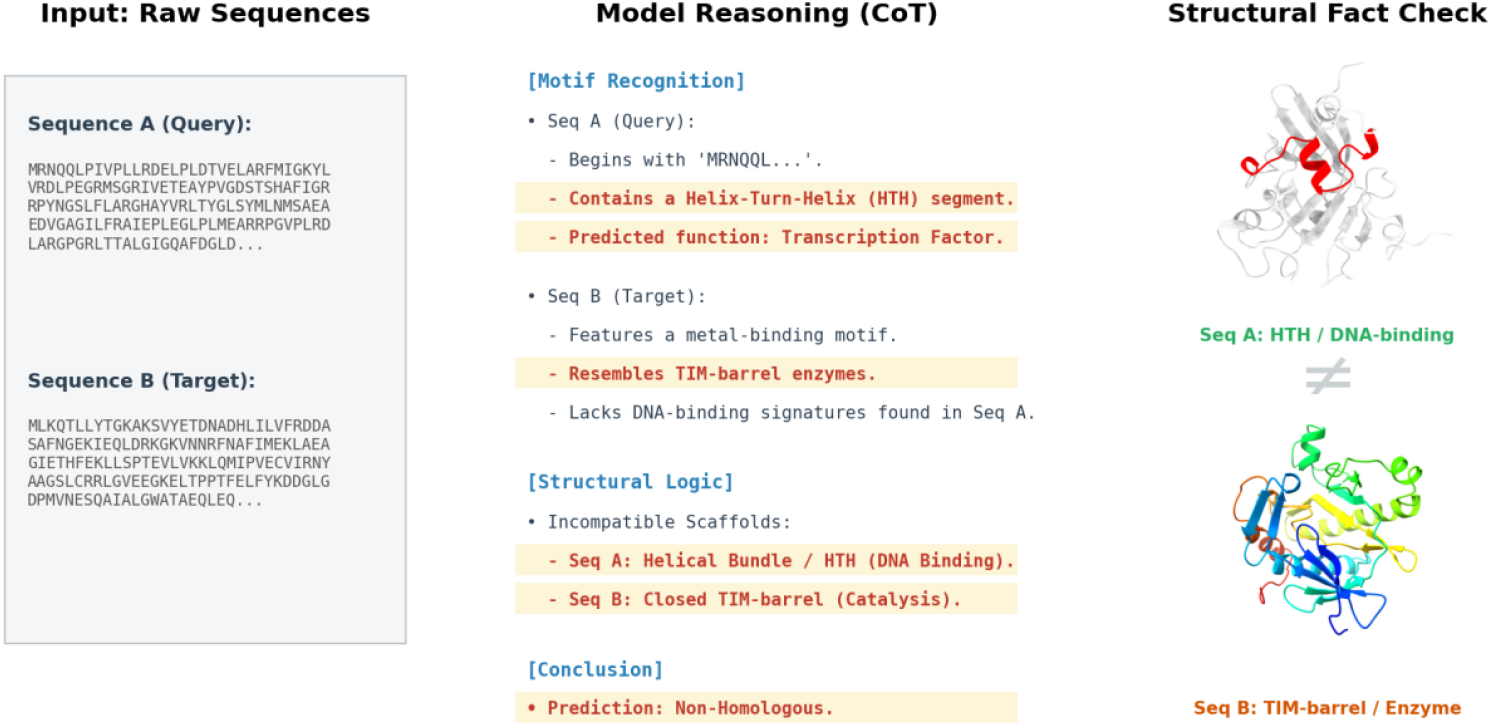
Exploratory: large models articulate structural reasoning. A representative Chain-of-Thought trace (Qwen-3) on a remote-homology pair: the model distinguishes a Helix-Turn-Helix motif from a TIM-barrel architecture and reaches a homology verdict without 3D coordinates. Shown as qualitative illustration only; because the model’s training corpus is undisclosed, this cannot be attributed to language-only transfer.

### Universality of Linguistic Transfer: From the Central Dogma to BioPAWS

Extending beyond proteins, PAWS-X-fine-tuned GPT-2 transfers to DNA homology detection above chance, and large models approach ceiling, indicating the transferable structural rules are modality-agnostic across nucleotides and amino acids. For Central Dogma mapping (verifying CDS-to-protein correspondence), purely linguistic fine-tuning is insufficient (near chance), whereas models with biological pre-training exposure succeed—indicating that syntax is transferable but the genetic “translation rules” require biological exposure. We consolidate protein/DNA homology, Central Dogma, and single-sequence property tasks (transcription-factor and solubility prediction) into the BioPAWS suite (Table 3), which distinguishes transferable syntactic tasks from semantic/functional tasks that remain hard.

**Table 3.** BioPAWS suite. GPT-2 (Base) denotes PAWS-X-only fine-tuning; GPT-2 (Bio) denotes pre-training exposure to biological data. Syntactic (homology) tasks transfer; the Central Dogma and functional tasks require biological exposure or large-scale reasoning.

| Category | Task | GPT-2 (Base) | GPT-2 (Bio) | Qwen-3 (zero-shot) |
| --- | --- | --- | --- | --- |
| Pairwise syntax | Protein homology | ~0.79 AUC | — | ~100% |
| Pairwise syntax | DNA homology | 78% | — | 100% |
| Cross-modal | Central Dogma (DNA→prot) | 55% (fail) | 81% | 83% |
| Semantic prop. | TF / solubility | — | — | ~65% |

## Discussion

### A partial isomorphism of universal syntax

Our controlled results show that a model trained only on English structure can perform protein-homology discrimination, implying that human language and the “language of life” share, at least partially, an underlying structural substrate. We frame this as a partial, sequence-similarity-level correspondence rather than a complete isomorphism: the transfer is real and pre-training-driven, but bounded.

### A cognitive-prior paradigm for AI-for-science, with known data

This challenges the assumption that biological problems require training from scratch on biological data: general models internalize abstract structural-comparison patterns usable as a prior. Crucially, by demonstrating the effect on GPT-2 and an English-only model we trained ourselves, we make “no biological training” a *verifiable* claim—a methodological standard we advocate for cross-domain transfer studies, given that large commercial models’ corpora cannot be audited.

### Limitations and the reasoning boundary

The transfer is discriminative and sequence-similarity level; it does not reach remote, fold-level homology, which defeats even specialized models under fine-tuning and appears to require reasoning that emerges only at large scale. Rather than contradicting the scaling observations, this delimits them: small models hit a representational ceiling, while large instruction-tuned models cross it via reasoning. Generative biological design and fold-level inference likely require hybrid systems pairing general reasoning with physics-based models. We view the present work as establishing, with verifiable controls, the existence and boundary of a genuine linguistic prior for biological structure.

## Data availability

Datasets—including exact file names, train/validation/test splits, and accession-level metadata—are released at https://huggingface.co/datasets/dnagpt/biopaws. All code for data construction, feature/shortcut analysis, finetuning, controls, the SmallWeb model, evaluation, and interpretability is available at https://github.com/maris205/biopaws.

### Supplementary S1: Prompt library for exploratory large-model evaluation

*Scope and caveat*. The templates below were used to query large third-party instruction-tuned models (e.g. Qwen-3, Gemini-class) in the *exploratory upper-bound* analysis only. Because the pretraining corpora of these commercial models are not public and may include biological data, results obtained with these prompts are reported as exploratory references rather than as controlled evidence; all controlled claims in the main text rest on GPT-2 and our English-only SmallWeb GPT-2, whose training data are fully known. The analogy language in the prompts (“paraphrase”, “mental folding”) reflects the wording given to the models and should not be read as a mechanistic claim. Inference used temperature=0.1, top_p=0.9 for deterministic decoding.

**S1.1 Protein / DNA homology detection (pairwise)**. *System*. “You are an expert bioinformatics assistant capable of linguistic transfer learning. Concept: an English sentence can be rearranged yet keep its meaning (paraphrase) or scrambled to lose its logic (adversarial). Analogy: homologous proteins/DNA are like paraphrases (different sequence, same structure/function); non-homologous or random sequences are like adversarial sentences (broken structural logic). Return strictly [{`“id”:1,”prediction”:”Homologous`”},…]; no explanations.” *User*. A JSON list of sequence pairs; the model judges each pair Homologous/Non-Homologous using “sequence syntax and structural integrity”.

**S1.2 Chain-of-thought reasoning (interpretability)**. *System*. “You are an expert structural biologist. Perform a ‘mental folding’ process: identify secondary-structure motifs and conserved domains, compare the two folding architectures, show step-by-step reasoning, then output `Prediction`: [`Homologous/Non-Homologous`].” Used to elicit abductive reasoning and motif identification (e.g. Helix-Turn-Helix, TIM-barrel) without 3D coordinates. Reported qualitatively.

**S1.3 Central-dogma mapping (DNA***→***protein)**. *System*. Framed as “abstract sequence alignment / cipher decryption”: the DNA is the source ciphertext, the protein the target plaintext; a coding relationship is a lossless translation, a non-coding one a glitch. The model judges Coding/Non-Coding from “information consistency and latent pattern alignment”, explicitly without codon tables.

**S1.4 Cluster-based single-sequence classification**. *System*. “Pattern-recognition engine”: given a mixed batch, separate sequences into two latent groups by shared sub-patterns/complexity and assign arbitrary but consistent labels*{*0, 1*}* Used for unsupervised-style DNA/protein binary tasks.

**S1.5 Few-shot in-context property prediction (e.g. solubility)**. *System*. Two reference examples (one per class) anchor a physicochemical decision (hydrophobic core vs. exposed patches); the model classifies a test batch by structural resemblance to the references.

**S1.6 Functional-element prediction (e.g. TF binding sites)**. *System*. Domain-knowledge-augmented prompt: identify whether DNA sequences carry motifs typical of transcription-factor binding sites (TATA box, zinc-finger motifs); output binary labels.

## Author contributions statement

L.W. conceived the project, designed and performed the experiments, analysed the data, and wrote the manuscript.

